# Decay of the CTCF paralog BORIS in neognathous birds

**DOI:** 10.1101/2025.02.12.637905

**Authors:** Katja Palitzsch, Thomas Wiehe, Peter Heger

## Abstract

BORIS (brother of the regulator of imprinted sites), the paralog of the genome organizer CTCF, originated at least 318 million years ago (Mya), in the ancestor of amniotes (mammals, reptiles, and birds). Based on results from chicken (*Gallus gallus*), the gene was thought to be absent from birds. Using comparative genomics of 59 bird species, we show that birds possess BORIS, but frequently experience severe degradation of the gene, as observed in *Gallus gallus*. The degradation events are restricted to neognathous birds, specific for the BORIS coding sequence, and occur multiple times independently on different branches. They comprise a wide range of molecular decay, from individual point mutations to the inactivation and/or loss of particular zinc fingers, to the almost complete disintegration of the gene. The decay is accompanied by relaxed evolutionary constraints on BORIS codons across neognathous birds and coincides with the accumulation of species-specific repetitive elements in degenerate loci. BORIS represents a case of a presently ongoing, convergent, and specific gene loss within a lineage. As possible explanation, we propose a link between the loss of BORIS and a shift in sperm and/or genital morphology during the evolution of Neognathae.

**Significance Statement:** The gene BORIS was believed to be absent from birds. However, genome analysis of 59 bird species reveals its presence, though it often underwent severe degradation in neognathous birds. These independent degradation events affect the BORIS coding sequence and range from point mutations to near complete gene disintegration. This decay correlates with relaxed evolutionary constraints and species-specific accumulation of repetitive elements. A potential link between BORIS loss and changes in sperm or genital morphology during neognathous bird evolution is suggested.

## Introduction

Isolated as a protein that binds to three regularly spaced CCCTC repeats upstream of the transcription start site, the CCCTC-binding factor CTCF was first described as transcriptional repressor of the c-*myc* oncogene in chicken [1]. In the meantime, it has become clear that CTCF is a highly conserved gene with very important roles in animal biology: As ubiquitously expressed DNA-binding protein with eleven zinc fingers (ZFs), CTCF is a key factor in 3D chromatin organization, genome partitioning, and gene regulation [2–6]. It operates through its unique ability to establish independent domains of gene expression known as topologically associating domains (TADs) [7, 8], a hallmark of cell-type specific gene expression and organismal development [9–14]. CTCF emerged in the ancestor of bilaterian animals, 540 million years ago. It is present in most bilaterians, such as insects, molluscs, and vertebrates, but absent from non-bilaterian animals and other eukaryotes [15, 16].

While CTCF is a single-copy gene in most bilaterians, including protostomes and non-vertebrate deuterostomes [16], some species and/or lineages experienced CTCF duplication. Prominent duplication events have been postulated in the ancestor of amniotes (mammals, reptiles, birds) and during early vertebrate evolution when CTCF gave rise to its paralog CTCFL (CTCF-like protein) or BORIS (brother of the regulator of imprinted sites) [17, 18]. BORIS is located within a synteny block highly conserved in amniotes. In humans, this synteny block is situated on chromosome 20 and spans seven genes, SPO11 (Meiotic recombination protein SPO11), RAE1 (RNA export 1 homolog), RBM38 (RNA binding motif protein 38), CTCFL (BORIS), PCK1 (Phosphoenolpyruvate carboxykinase 1), ZBP1 (Z-DNA binding protein 1), and PMEPA1 (Prostate transmembrane protein, androgen induced 1) [19]. The direct neighbours of BORIS, RBM38 and PCK1, are ancient genes and regulate gene expression at the post-transcriptional level (RBM38: [20, 21]) or perform basic metabolic tasks (PCK1: [22, 23]). In contrast, CTCF’s neighbour genes, RIPOR1 (RHO family interacting cell polarization regulator 1) and CARMIL2 (capping protein regulator and myosin 1 linker 2), exhibit highly dissimilar sequence signatures and molecular functions [24, 25], suggesting that CTCF duplicated and transposed to its new genomic location as an isolated gene.

As a result of their evolutionary history, CTCF and BORIS share a high degree of homology in their central zinc finger region. This part of the protein is characterized by 74 % amino acid identity, similar DNA binding properties, and a conserved genomic structure [17, 26–28]. In both paralogs, the ZF region is composed of seven exons, and each of these carries information for one and a half, or two (exons 3, 7, 8), zinc fingers [17, 26]. Despite these similarities, BORIS and CTCF differ substantially in their *N* - and *C*-terminal domains [26, 29]. As a consequence, the two proteins interact with different partner proteins and have distinct developmental functions and expression patterns [17, 26, 30, 31]. While CTCF is ubiquitously expressed from early development on [32–37], BORIS expression in mammals seems to be restricted to germline cells [17, 26, 38] where it regulates genes with a role in spermatogenesis and spermatid differentiation [39–41]. Due to its function as a transcriptional regulator of critical genes, amplification of the BORIS locus and/or misexpression of BORIS target genes are involved in the etiology of a number of cancers. According to its testis-specific expression and the irregular expression in cancer tissues, BORIS is classified as a member of the cancer-testis (CT) genes [for review, see: 42, 43].

With the growing amount of genomic data from a wide variety of organisms throughout all kingdoms of life [for review, see 44], evidence has accumulated that gene loss is a powerful evolutionary force, similar to evolution by gene duplication [45]. There are numerous examples of gene loss throughout the animal kingdom, including the regression of vision and pigmentation in Mexican cavefish [46–48], the loss of vitamin C synthesis in primates and other vertebrates [49–51], or the loss of developmental regulators such as Hox genes and CTCF in nematodes [52, 53]. In particular, there is increasing evidence that species that share a similar ecological niche experience the loss of the same gene(s) in a convergent manner. Systematic computational screens revealed that, for example, carnivorous and herbivorous mammals, animals with low visual acuity, or aquatic mammals encounter convergent loss of the same genes independently [54–56]. Expanding on these studies, we describe here numerous mutational states that characterize the ongoing loss of a putative gene regulatory protein, BORIS, from a distinct lineage of birds. We support our main findings by analyses of repeat element content and selective forces in the BORIS locus and conclude that this locus is destined for extinction in neognathous birds.

## Results

### Absence of a functional BORIS gene in *Gallus gallus*

The CTCF paralog BORIS, also known as CTCF-like (CTCFL) [26], originated in the ancestor of amniotes 318 Mya [17] and therefore is expected to exist in the genomes of reptiles, mammals, and birds. However, apart from a small fragment with similarity to zinc finger one (ZF I) [17], previous studies failed to detect BORIS in the chicken genome despite its presence in other bird and amniote species [19]. To resolve the conflicting findings, we investigated by a combined *in vitro* and *in silico* approach if there is a functional BORIS gene in the *Gallus gallus* genome. Utilizing computational searches with a hidden Markov model for the BORIS/CTCFL coding sequence, we identified the 36 AA CTCFL fragment described by Hore et al. [17] and three additional genomic fragments with similarity to BORIS/CTCFL in the *Gallus gallus* genome (Figure 1; Table 1).

**Figure 1:**
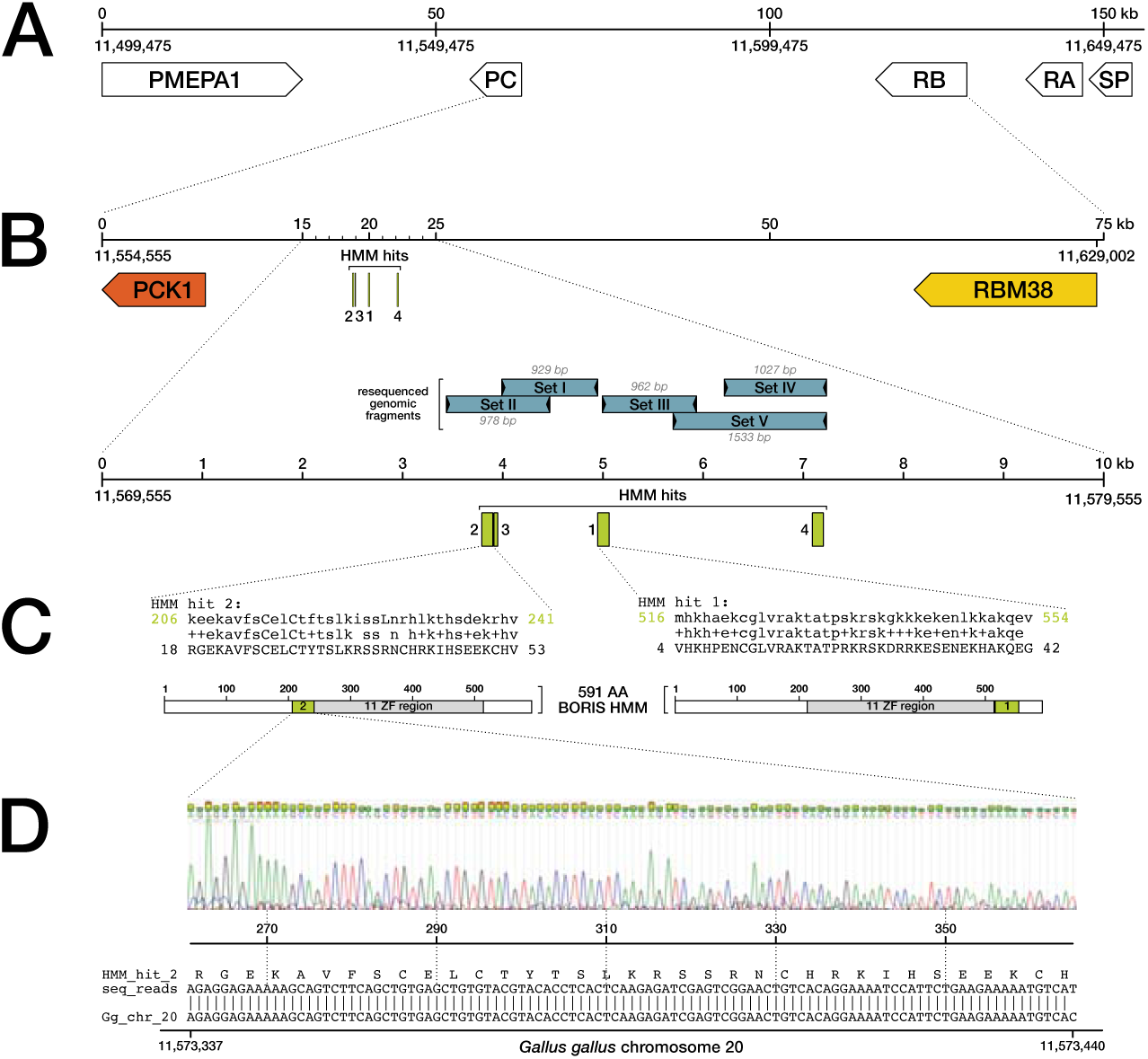
Presence of non-functional BORIS fragments within the *Gallus gallus* BORIS syntenic region. **A:** Syntenic block of the five genes PMEPA1, PCK1 (PC), RBM38 (RB), RAE1 (RA), and SPO11 (SP), surrounding the BORIS/CTCFL locus on *Gallus gallus* chromosome 20 (position 11 499 475 to 11 653 697 in genome assembly version GRCg6a). Order and arrangement of the genes are conserved in amniotes [19]. Gene sizes and distances are drawn to scale, gene orientation is indicated by arrowheads. **B:** The 75 kb region between the BORIS neighbour genes PCK1 and RBM38 with four BORIS fragments identified by HMM searches (green; HMM hits 1–4) and five re-sequenced genomic regions, covering a 3.7 kb area (blue Set I to Set V; for PCR/sequencing primers, see Table 2). HMM hits are numbered according to Table 1. Drawn to scale. **C:** Alignment details of HMM hits 1 and 2, as reported by HMMSEARCH [57]. The identified genomic fragments (lower sequence; position on ORF indicated by numbers) are aligned to a 591 AA hidden Markov model of BORIS (upper sequence, match on HMM indicated by green numbers). Conserved residues between identified ORF and BORIS HMM are reported in the central line. Corresponding hit regions within the BORIS HMM are marked in green in the cartoon below. ZF region: zinc finger region of BORIS. **D:** Exemplary sequencing detail: Chromatogram of a Set II sequencing read across HMM hit 2 (top) and alignment between the obtained sequence/ORF and the corresponding genomic region on *G. gallus* chromosome 20 (bottom).

**Table 1:**
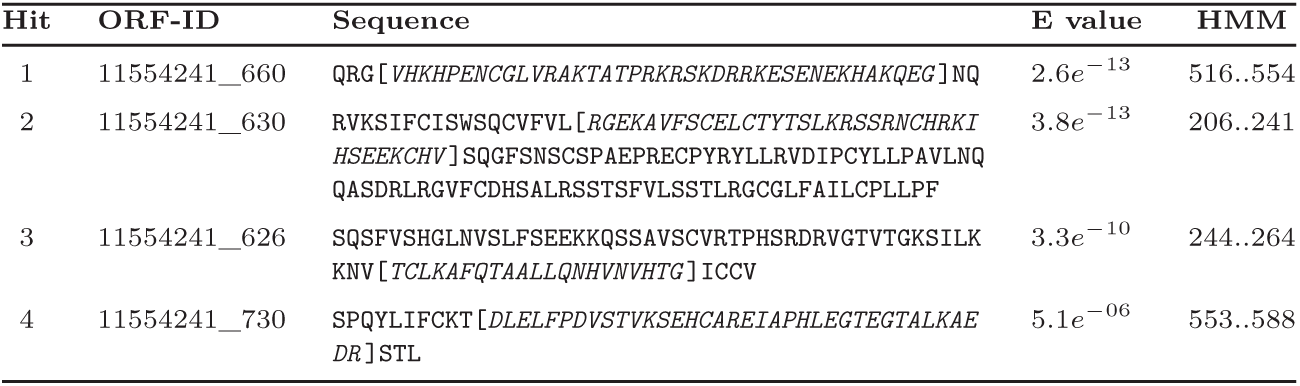
BORIS fragments within the Gallus gallus PCK1–RBM38 syntenic region. Hit number and ORF-ID of four *Gallus gallus* ORFs with similarity to BORIS, as revealed by HMM searches (columns one and two). ORFs are derived from chromosome 20 of the *Gallus gallus* genome assembly, version GRCg6a (NC_006107.5, contig 11628597). Column three displays the corresponding ORF sequence. The actual HMM search hit region, corresponding to a BORIS fragment, is positioned within square brackets and highlighted in italics. Rightmost columns indicate the region of the 591 AA BORIS HMM to which the detected fragments display similarity, and the corresponding similarity measure (E value). For graphical representation, see Figure 1.

**Table 2:**
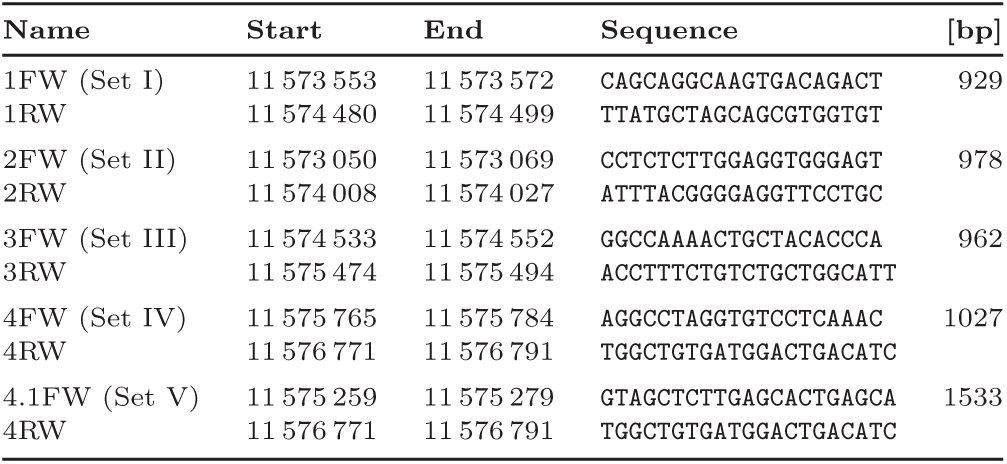
Primer list for the PCR-based amplification of genomic DNA fragments. Columns two and three show start and end coordinates of the respective primer on chromosome 20 of the *G. gallus* genome assembly, version GRCg6a. Column five indicates the expected amplicon size for the respective primers.

As expected from synteny information in other amniotes [19], all four ORFs were located between the genes PCK1 and RBM38 where the BORIS locus is situated.

The pieces corresponded to ZF I (HMM hit 2, as identified by Hore et al. [17]), the majority of ZF II (HMM hit 3), and a conserved protein sequence directly after ZF XI, separated into two exons (HMM hits 1 and 4). Together, they comprised 128 AA or 19.3 % of BORIS’ expected size (663 AA in humans) and spanned ∼ 3.4 kb in the genome, with no identifiable traces of the missing parts of the gene in the corresponding genomic region. To confirm that the degenerate configuration derived *in silico* reflects the situation *in vivo*, we PCR-amplified and sequenced from chicken genomic DNA a 3.7 kb region extending from the detected ZF I fragment (HMM hit 2) to the conserved region after ZF XI (HMM hit 4; Figure 1; Table 1). We would expect to find remnants of the highly characteristic CTCF/CTCFL zinc finger domains [15] in the amplified DNA if the gene was present in *Gallus gallus*. However, the re-sequenced DNA closely matched the genome assembly and thus did not contain additional parts of BORIS missing from the assembly (Figure 1D). In *G. gallus*, the region from the start of HMM hit 2 to the end of HMM hit 1 corresponds to exons 3 and 9 (AA 206– 554; see Figure 1C and Table 1) of the 591 AA BORIS HMM and covers 1277 bp in the genome assembly. In contrast, the corresponding region (BORIS AA 206–554) of the closely related galliform *Numida meleagris* extends over seven exons and 6136 bp, suggesting that several central exons of BORIS are missing in *G. gallus*. Together, these data confirm the correctness of the *Gallus gallus* genome assembly at the BORIS locus and demonstrate the absence of a functional BORIS gene in this species despite the retention of four exon fragments in the original syntenic context.

### The degeneration of BORIS is recent, recurrent, and restricted to neognathous birds

To determine if BORIS is also prone to degenerative events in birds other than *Gallus gallus*, we compiled a comprehensive dataset of 59 bird genomes, including Paleognathae and other Galliformes (Supplementary Tables S1, S2), and examined in detail the state of the BORIS gene in these species. As for *Gallus*, we extracted from the genomes the syntenic region surrounding the BORIS locus, from the genes PCK1 to RBM38. We then translated the corresponding sequences into six reading frames and scanned the resulting ORFs with a 591 AA BORIS hidden Markov model (HMM) to detect with high specificity ORFs similar to the BORIS protein, independently of potentially missing or erroneous genome annotations. With the help of this method, we identified and annotated across all bird species ORFs of BORIS’ highly conserved zinc finger region. Our results show that the BORIS zinc finger region is complete and intact in the eleven paleognathous species of our collection, spanning all known families of this monophyletic clade [Prum et al. [58]; Figure 2]. In contrast, we detected a large number of degenerative events in the BORIS genes of neognathous birds (Figure 2). Mapped onto a bird phylogeny, our results revealed that (i) BORIS mutations are detectable only in neognathous birds but are absent from Paleognathae; (ii) changes in BORIS comprise a wide spectrum of molecular decay: from point mutations affecting zinc-complexing residues—and therefore DNA binding capabilities [59]—(e. g. in *Columba livia*, *Amazona aestiva*, *Nipponia nippon*) to frame shifts and stop codons (e. g. in *Nestor notabilis*, *Tauraco erythrolophus*, *Calidris pugnax*), to the loss of individual zinc fingers or exons (e. g. in Passeriformes, *Cariama cristata*, or *Cuculus canorus*), to the severe disintegration of the entire genomic locus, as in *Acanthisitta chloris*, *Dryobates pubescens*, or *Gallus gallus*; (iii) different bird lineages acquired distinct modifications of the BORIS locus, on a global scale across the dataset as well as within monophyletic clades (e. g. Columbaves) or between sister species (e. g. *Dryobates pubescens* vs. *Merops nubicus*); (iv) mutations preferentially affect the *C*-terminal part of the BORIS zinc finger region (30 species with mutations in ZF VII–XI) while changes in the N-terminal part are less prevalent (14 species with mutations in ZF I–V; Figure 2). Although, among the birds in our set, the *G. gallus* BORIS locus is most severely affected by degeneration, substantial damage to BORIS is also observed in several other species (e. g. *Acanthisitta chloris* or *Dryobates pubescens*). Together, we identified 26 independent BORIS mutational profiles in our tree of 48 neognathous species, and therefore mutations in every other species (54.2 %; Figure 2).

**Figure 2:**
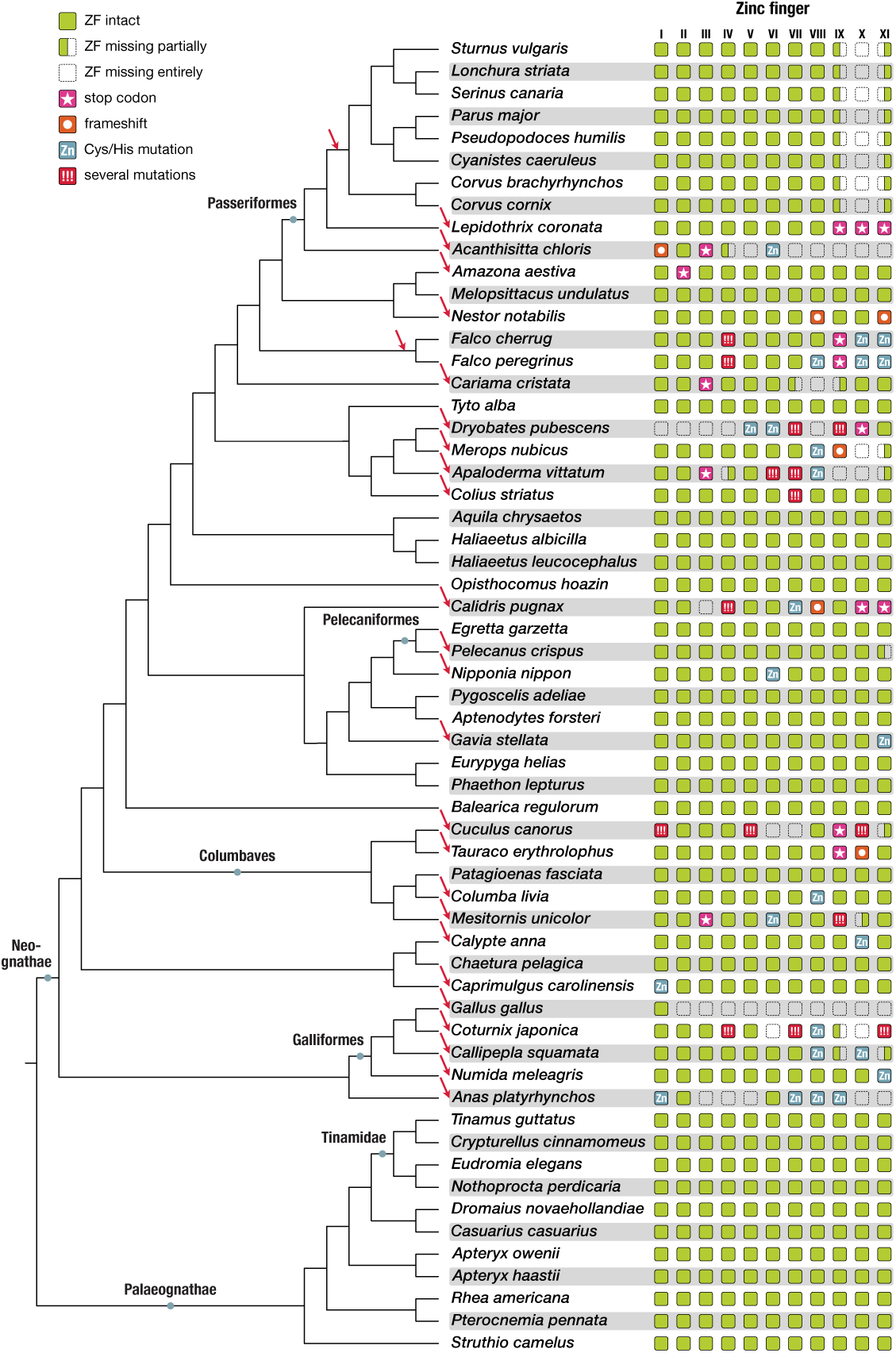
Convergent degeneration of CTCFL/BORIS in neognathous birds. **Left:** Cladogram depicting the relationships of 59 bird species. Tree topology after https://phylotastic.org/ and Prum et al. [58], Wang et al. [60], Weir et al. [61]. Major bird lineages are indicated by grey dots. **Right:** Presence and mutational state of the eleven CTCFL/BORIS C_2_H_2_ zinc fingers (ZF I–XI), mapped onto the bird phylogeny (left). Intact and complete zinc fingers are depicted as green squares, damaged zinc fingers are classified by icons (see legend at top left). Red arrows indicate independent degeneration events within a given lineage.

To investigate whether degradation events of the BORIS gene are detectable in other amniotes as well, we used the same work flow (scanning ORFs of the BORIS syntenic region with our BORIS HMM) and analyzed the mutational state of BORIS in representative sets of the two other amniote clades, in mammals and reptiles. The corresponding results clearly show that BORIS is intact in all investigated mammalian (*n* = 38) and reptilian (*n* = 16) species (Supplementary Figures S1,S2) and confirm that specifically neognathous birds experienced a loss of this gene, as its integrity in paleognathous birds had suggested (Figure 2).

### Degenerative events are specific for the BORIS locus

If the observed losses were a result of chromosomal rearrangements around the BORIS locus in Neognathae, we could expect that BORIS’ neighbour genes suffer from such modifications as well. To test this possibility, we investigated the completeness and integrity of the two BORIS flanking genes, PCK1 and RBM38, across the bird phylogeny. As proxy for intactness we selected the two genes’ coding sequence and subsequently collected these from the proteomes of different bird species by BLAST searches at NCBI. We then created multiple sequence alignments of the two proteins (PCK1 and RBM38), each consisting of sequences from at least 40 species. Finally, we inferred from these alignments the mutational state of the two neighbour genes across birds.

Neighbour gene RBM38 (RNA-binding protein 38) is a 215 AA RNA-binding protein with a role in transcript stabilization. It is highly conserved in birds and other species. Regardless of mutations at the BORIS locus, the RBM38 gene is complete and undamaged in all examined birds (Supplementary File 1). Similarly, alignments of PCK1 (Cytosolic phosphoenolpyruvate carboxykinase 1), the rate-limiting enzyme in gluconeogenesis (622 AA in birds), show a strong conservation of its protein sequence across birds, without signs of damage or deletion (Supplementary File 2). In particular, species with defects in BORIS do maintain intact neighbour genes RBM38 and PCK1, e. g. *Gallus*, *Acanthisitta*, or *Dryobates*. These observations indicate that the degenerative events observed in neognathous birds are restricted to and specifically affect the BORIS locus.

### Accumulation of species-specific repetitive elements in degenerate BORIS loci

Genomic regions free of coding sequence or regulatory function may evolve with little selective constraint and accumulate nucleotide substitutions, insertions, or deletions at a faster rate than sections under purifying selection. In particular, unconstrained regions may tolerate the insertion of repetitive elements.

To test whether degenerate BORIS loci are susceptible to repeat insertion, we carried out a thorough analysis of repeat element abundance in 18 bird species with intact and degenerate BORIS loci (Supplementary Table S3) by combining a de novo and a library-based approach for repeat detection. First, we generated a comprehensive bird de novo repeat library from the joined output of independent RepeatModeler (detection of transposable elements, tandem repeats, and LTRs) and Mitetracker (detection of MITEs—miniature inverted-repeats) runs on all 18 genomes. We then carried out RepeatMasker analyses on all genomes, using the combined de novo library. In addition, a second RepeatMasker run on all 18 genomes utilized a standard RepeatMasker library (RepBase Release 20181026) for repeat detection (results are summarized in Supplementary File 3).

In agreement with previous studies [62, 63], we find that several repeat classes are highly abundant in birds on a genome-wide scale (SINEs, LINEs, CR1 and LTR elements, DNA transposons, Small RNA) while others could not be detected at all (CRE/SLACS, R1/LOA/Jockey, BEL/Pao, Ty1/Copia, En-Spm, or PiggyBac elements; Supplementary Figure S3). For most species, the RepeatMasker library-based pipeline reveals more repetitive elements in total than does the de novo pipeline (Supplementary Figure S3). A few repeat classes, however, could only be detected by the de novo library (e. g. the R2/R4 class, «Other», and Simple Repeats; Supplementary Figure S4), demonstrating the value of a two-tiered strategy for repeat identification. Second, we find that highly abundant retroelements, in particular the classes «LINE», «CR1», «LTR element», and «retroviral element», are more prevalent in neognathous birds than in paleognathous species (Supplementary Figure S4), supporting the view that the evolution of Neognathae is accompanied by an expansion of retroelements that utilize RNA intermediates. In contrast, DNA-based repeat elements are more prevalent in paleognathous birds despite their generally lower abundance (Penelope, DNA transposons, Tourist/Harbinger, Simple Repeats; Supplementary Figure S4), suggesting that Neognathae and Paleognathae differ in their mechanisms of repetitive element control. *Dryobates’* unusually high repeat content is in line with previous findings [63], demonstrating the robustness of our repeat detection pipeline (Supplementary Figure S3).

Next, we analysed in detail the repeat landscape within the BORIS syntenic region of the 18 bird species. From species with intact BORIS coding region we deduced that complete BORIS loci extend over ∼13 kb in birds. We then identified BORIS marker exons in all bird genomes and aligned the 13 kb region accordingly, thereby obtaining genomic coordinates of the supposed consensus BORIS locus. When we looked for repeat elements present within these coordinates, we found that both, repeat counts per kb of genomic sequence and total repeat counts, were elevated in degenerate BORIS loci of neognathous birds, as compared to intact loci (Figure 3A,B), although statistical tests with a significance threshold of *p* = 0.05 narrowly failed to recover significant differences (Mann-Whitney U test: statistic: 12.00, *P* value = 0.09; Kruskal-Wallis test: H statistic: 3.125, *P* value = 0.077). In contrast, repeat contents of intact BORIS loci were almost identical between the two bird groups (Mann-Whitney U test: statistic: 15.00, *P* value = 0.86; Kruskal-Wallis test: H statistic: 0.077, *P* value = 0.782), implying a true, albeit not significant difference in repeat counts between intact and degenerate BORIS loci. Of 90 distinct repeat elements that populate the BORIS loci of the 18 analysed species, 68 (75.6 %) are present only once in a single locus of a single species, 18 elements (20.0 %) occur in two or three copies (often within the same locus), and four (4.4 %) elements occur 5 to 8 times (Supplementary File 4). The most abundant repetitive elements (5 to 8 counts) are MER131 («medium reiterated frequency repeat» 131), a conserved interspersed repeat common in Euteleostomi [64]); MIR1_Amn («mammalian-wide interspersed repeats»), an ancient family of tRNA-derived SINEs (short interspersed nuclear elements) [65]; and two repeats detected by our de novo library (Drypub_rnd-1_family-156 and Strtur_ltr-1_family-80; Supplementary File 4). According to their conservation across species, all these elements share a long history and originated in the ancestors of birds or earlier. On the other hand, most elements that occur exactly once are direct repeats (DRs; 45 of 68 unique elements or 66.2 %), in particular CR1 repeats common in neognathous birds (Supplementary Figure S4) and other amniotes [66]. Detailed analyses reveal that all direct repeats within BORIS loci are restricted to a single species (Supplementary File 4), suggesting that they emerged in their host and are evolutionarily young compared to the more ancient elements mentioned above. When we looked at the abundance of ancient and young repeats, we found that the latter are significantly overrepresented in degenerate BORIS loci (Kruskal-Wallis test: H statistic: 4.109, *P* value = 0.043; Figure 3C,D). In contrast, the number of ancient repeats is not significantly different in intact and degenerate BORIS loci (Kruskal-Wallis test: H statistic: 0.433, *P* value = 0.510). Thus, degenerate BORIS loci emerged independently in closely related neognathous birds (Figure 2) and contain a large number of evolutionarily young repetitive sequences compared to intact loci (Figure 3), arguing that BORIS degeneration and repeated invasion of its locus are species-specific processes that take place simultaneously and possibly influence each other.

**Figure 3:**
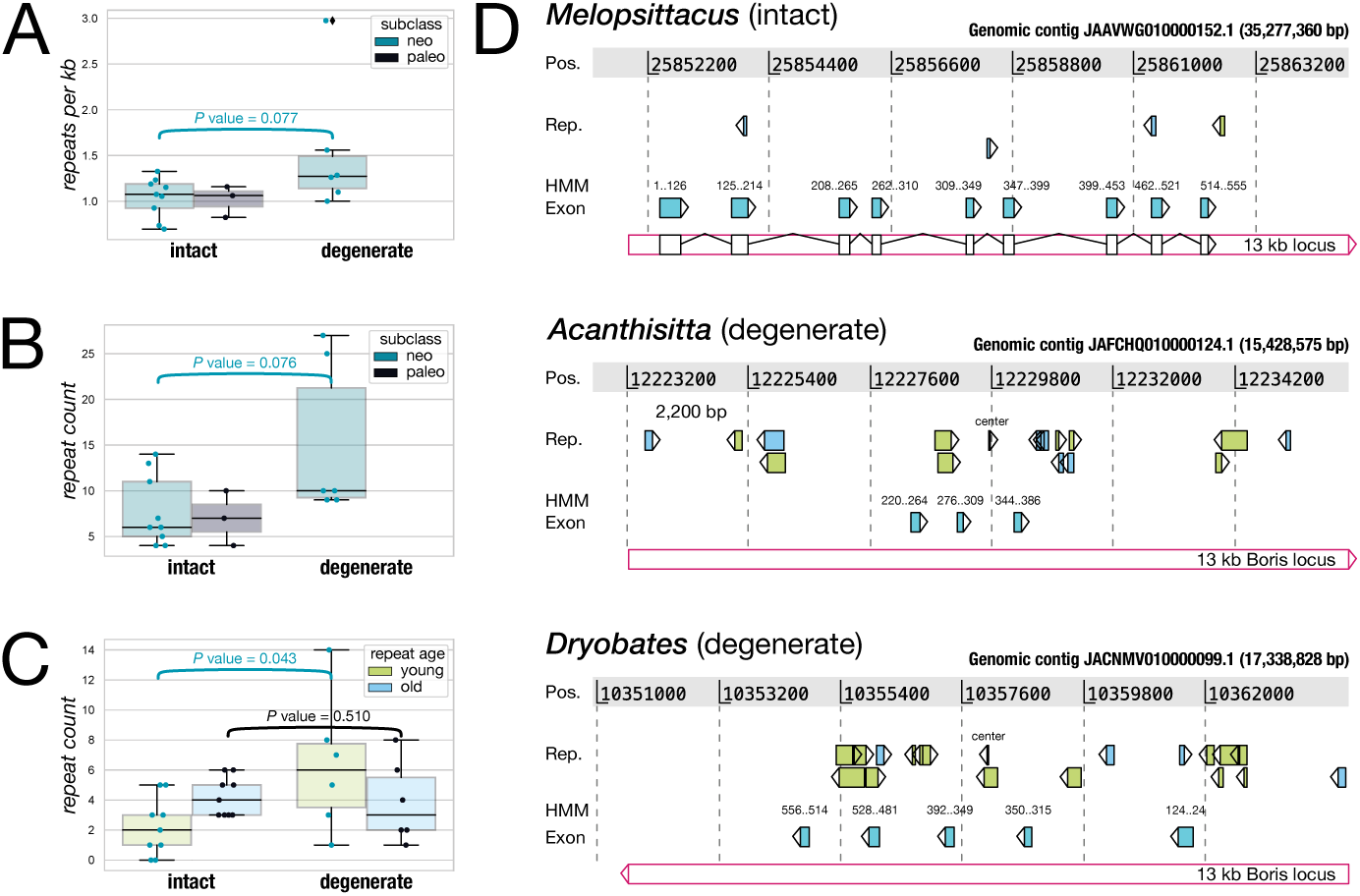
Accumulation of repetitive elements in degenerate BORIS loci. **A:** Box-plots showing the total number of repeat elements per kb detected on the genomic contig carrying BORIS, in species with intact (left; n=12) and degenerate BORIS loci (right; n=6: *Acanthisitta chloris*, *Anas platyrhynchos*, *Calidris pugnax*, *Cuculus canorus*, *Dryobates pubescens*, *Gallus gallus)*. **B:** Boxplots showing the number of repeat elements detected within the 13 kb BORIS locus in species with intact (left) and degenerate (right) BORIS. **C:** Boxplots showing the number of ancient and young (species-specific) repeat elements in intact vs. degenerate BORIS loci. **D:** Representative intact and degenerate BORIS loci of three bird species, annotated with repeat elements (ancient: light-blue, young/species-specific: yellow) and BORIS exons (cyan). Genomic contigs carrying the respective BORIS locus are referenced at the top right. Four labels at the left indicate (i) the genomic position of an element on the respective contig («Pos.», highlighted as grey box), (ii) the location and orientation of repetitive elements («Rep.»), (iii) the region on the 591 AA BORIS HMM where exons matched («HMM»), and (iv) the location and orientation of BORIS exons as detected by HMM search («Exon»). Red boxes at the bottom display the 13 kb BORIS consensus region. Drawn to scale.

### Relaxed selection on BORIS codons in neognathous birds

The recurrent degeneration of BORIS in neognathous birds indicates that the gene might become dispensable in this bird lineage. We should therefore be able to observe a difference in the selective pressure on BORIS between Neognathae and Paleognathae, across which BORIS is consistently well conserved (Figure 2). As a measure of selective constraint we examined the nonsynonymous vs. synonymous substitution rate (*dN/dS*) in the two bird lineages. To this end we created a codon-based multiple sequence alignment of the BORIS zinc finger region across the avian phylogeny. We included in the alignment ten paleognathous and 19 neognathous birds with intact, full-length coding sequence, but excluded species with degenerate BORIS. Then, we determined the *dN/dS* ratios in pairwise comparisons of all possible sequence combinations using the maximum likelihood approach implemented in CODEML [67]. We found that comparisons within paleognathous species (PP) had a low *dN/dS* ratio (0.001 to 0.25 in 45 unique comparisons; mean: 0.069), as expected for a conserved gene under functional constraint. On the other hand, *dN/dS* ratios between neognathous birds (NN: 0.099 to 0.699 in 171 comparisons; mean: 0.204) and between neo- and paleognathous species (NP: 0.069 to 0.35 in 190 comparisons; mean: 0.165) were markedly higher, suggesting a relaxation of selective constraint by higher nonsynonymous substitution rates in the neognathous lineage (Figure 4A). When we calculated *dN/dS* ratios in an analogous way for two control genes, PCK1 (18 neognathous, five paleognathous species) and CTCF (18 neognathous, eight paleognathous species), we observed that their *dN/dS* ratios were uniformly low across the bird tree, i. e. across NN, NP, and PP comparisons (Figure 4B). Statistical tests underscore these results: while there is no significant difference between NN and PP comparisons for the genes PCK1 (H statistic: 0.507; *P* value = 0.476) and CTCF (H statistic: 3.428; *P* value = 0.064), *dN/dS* values for the BORIS coding sequence are significantly higher in NN comparisons than in PP comparisons (H statistic: 210.478; *P* value = 1.08*e*-47), as determined by nonparametric Kruskal-Wallis tests (Figure 4B). Thus, BORIS is maintained as a functional gene under purifying selection in Paleognathae. In contrast, functional constraint is relaxed (higher *dN/dS* values) specifically in neognathous birds, even in those that still possess an intact BORIS zinc finger domain.

**Figure 4:**
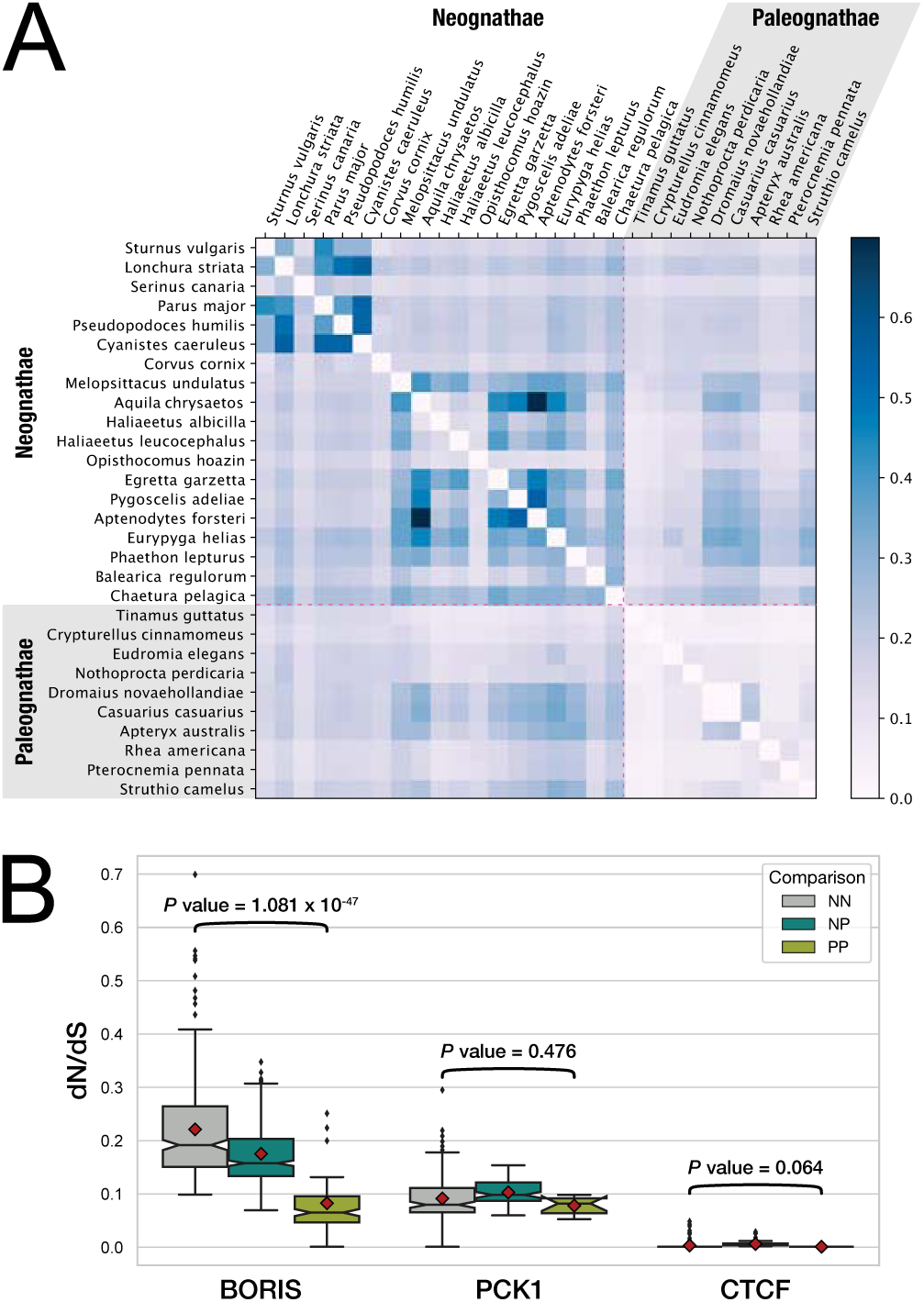
Maximum likelihood analysis of BORIS codon evolution in neognathous and paleognathous birds. **A:** Heatmap of *dN/dS* ratios derived from pairwise sequence comparisons of ten paleognathous and 19 neognathous bird species. Paleognathous birds are highlighted by a grey background and a pink dashed line. *dN/dS* ratios were determined by CODEML [67] using the ZF coding region of intact BORIS genes. Note that *dN/dS* values are plotted in logarithmic scaling. **B:** Boxplot summaries of *dN/dS* ratios for BORIS (same data as in heatmap above) and two control genes, PCK1 and CTCF, on the basis of pairwise comparisons between neognathous and paleognathous species (neognathneognath = NN, grey; neognath-paleognath = NP, bluegreen; paleognath-paleognath = PP, yellowgreen). Red diamonds denote the mean of a dataset. Note that *dN/dS* results for the ZF region of the BORIS paralog CTCF are extremely low and uniform in all analysed comparisons, preventing the formation of a box. *P*-values < 0.05 indicate a significant difference between *dN/dS* values derived from NN vs. those derived from PP pairwise comparisons (Kruskal-Wallis nonparametric tests).

While these findings establish the ongoing relaxation of selective pressure on BORIS across Neognathae, they do not allow to dissect the relative contributions of individual phylogenetic branches to the overall signal. Also, pairwise comparisons are not independent from each other and may be distorted by an extreme inflation of the *dN/dS* ratio in only one, or a few, branches [68]. To obtain a more detailed picture, we mapped the substitution events inferred by CODEML onto a consensus bird phylogeny, counting nonsynonymous and synonymous substitutions per branch. In paleognathous birds, we find on most internal branches (20 out of 21) much lower counts of non-synonymous than synonymous substitutions. Only on the branch leading to the genus Apteryx the counts are similar to each other. In contrast, within Neognathae, non-synonymous counts are similar to or even larger than synonymous ones on 16 out of 37 branches (Figure 5). Importantly, we find such an excess of nonsynonymous substitutions across the entire neognathous avian tree, suggesting that relaxed constraint on BORIS appears to be a general phenomenon in these birds and not the result of few, particularly fast evolving species that might bias the overall view. Still, one extreme signal with many more nonsynonymous than synonymous substitutions can be detected on the branch leading to Psittacopasserae (parrots and passerines [69]), indicating that substantial changes in the BORIS coding region occurred during the evolution of this monophyletic clade (Figure 5). This may point to a phase of accelerated evolution in birds around 60 Mya and is in line with difficulties to resolve the phylogenetic placement of passerines [69].

**Figure 5:**
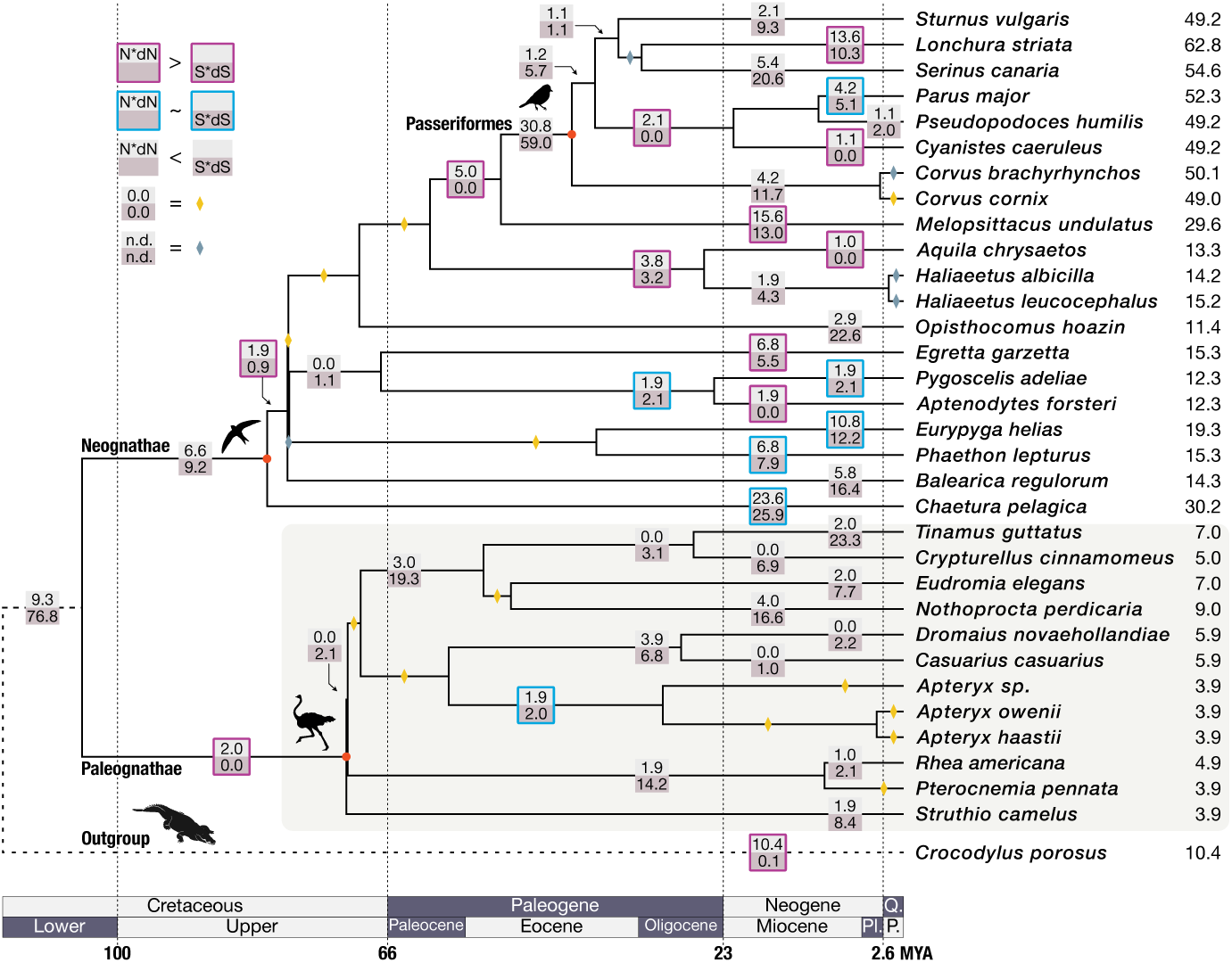
Branch-specific mapping of BORIS codon evolution in birds. Branch-specific mapping of substitution events onto a consensus phylogeny of 12 paleognathous birds, 20 neognathous birds, and the outgroup species *Crocodylus porosus*. Tree topology and time scale are derived from **Error! Hyperlink reference not valid.** Species silhouettes for major clades (red dots) were downloaded from https://www.phylopic.org/. Branch numbers indicate *N* ×*dN* (upper) and *S*×*dS* (lower) values as obtained from CODEML [67], using the codon-aligned ZF region of intact BORIS genes as input. Pink frames highlight branches where *N* × *dN > S* × *dS*. Blue frames indicate branches with *N* × *dN* ∼ *S* × *dS*. Yellow diamonds identify branches with *N* × *dN* = *S* × *dS* = 0.0. Grey diamonds label branches with undetermined *N* × *dN*, *S* × *dS* values. Paleognathous birds are highlighted by a grey background. Pl: Pliocene, P: Pleistocene, Q: Quaternary.

### Altered sperm morphology in neognathous birds

In mammals, BORIS is expressed in testes and is associated with sperm development and differentiation [39–41]. In contrast, reptiles express BORIS in gonads *and* in somatic tissues according to a comparative study [17]. So far, nothing is known about expression patterns and BORIS functionality in birds. Assuming that BORIS is also important for sperm development in birds and reptiles, as is in mammals, sperm from Neognathae may differ from sperm from Paleognathae/Reptilia, reflecting the molecular difference of BORIS conservation between Neognathae and Paleognathae on a phenotypic level. To test this idea, we extracted measurements from the Sperm Morphology Database [70] on the length of sperm heads, mid pieces, and flagella for Aves and Reptilia. Since the database provides only two datasets for paleognathous birds, we combined this group and Reptilia to a larger dataset and compared it to the Neognathae (Figure 6).

**Figure 6:**
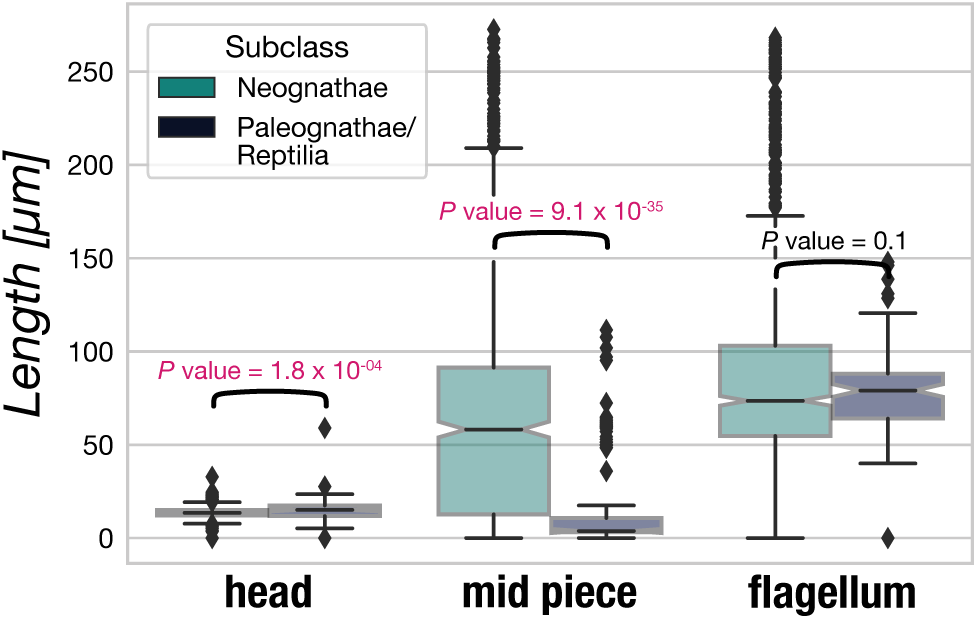
Comparison of morphological characteristics of sperm from Neognathae and Paleognathae/Reptilia. The lengths of sperm heads (left), mid pieces (middle), and flagella (right) from neognathous birds (554 species) are compared to the corresponding data of paleognathous birds and reptiles (122 species). All data are taken from the SpermTree Database (https://spermtree.org/ [70]). *P*-values *<* 0.05 are highlighted in red and indicate that a morphological trait differs significantly between Neognathae and Paleognathae/Reptilia, as determined by Kruskal-Wallis tests.

Our comparisons reveal that sperm head lengths differ slightly between Neognathae and Paleognathae/Reptilia (Neognathae: 14.13±6.49 μm; Paleognathae/Reptilia: 15.41 ± 5.53 μm; Figure 6), as do flagellum lengths (Neognathae: 101.86 ± 58.77 μm; Paleognathae/Reptilia: 79.23 ± 19.49 μm; Figure 6). In contrast, there are significant differences in the length of mid pieces between the two groups. The mid pieces of sperm from Neognathae are clearly larger (81.92 ± 66.99 μm) than those of Paleognathae/Reptilia (18.53 ± 27.75 μm; Figure 6). This is confirmed by non-parametric statistical tests (Kruskal-Wallis) that recover significant differences in the length of sperm heads and mid pieces between Neognathae and Paleognathae/Reptilia (head: H statistic: 13.997, *P*-value = 1.83*e*-04; mid piece: H statistic: 151.285, *P*-value = 9.08*e*-35; flagellum: H statistic: 2.462, *P*-value = 0.116).

Due to the low abundance of paleognathous species, the difference in sperm mid piece length reported here essentially reflects a difference between Neognathae (554 data points) and Reptilia (120 data points; compare Figure 6 to Supplementary Figure S5A). However, additional comparisons between Paleognathae and Reptilia and between Paleognathae and Neognathae corroborate that mid piece and flagellum lengths of Paleognathae are well below the interquartile range of neognaths and reptiles (Supplementary Figure S5B, C), suggesting that the two branches of birds indeed show a significant difference in the length of their sperm mid pieces.

These findings indicate that a relaxed selective pressure on BORIS and its tendency to degrade are correlated with an increase in the length of sperm heads and mid pieces in Neognathae. Whether these alterations of sperm morphology in neognathous birds are caused by a change in BORIS function or expression needs to be determined in future experiments.

## Discussion

Here we report that the BORIS gene, a paralog of the transcription factor CTCF, is conserved in paleognathous birds, but is degrading in many species of its sister clade, the neognathous birds. Loss of BORIS cannot be explained by a single event in the neognathous ancestor. Instead, our comparison of genomic sequences suggests a series of independent mutational damages since the Paleognathae and Neognathae split about 108 Mya. We were able to identify traces of the BORIS gene in the genomes of all analyzed Neognathae and found no case of complete absence. However, we discovered a wide range of partial losses, with damage from mild to severe. We conclude from these findings that the degradation of BORIS is a recurrent and still ongoing process. Although the BORIS gene has apparently remained intact in some Neognathae, our analysis of nonsynonymous vs. synonymous substitutions suggests that purifying selection is reduced in Neognathae compared to Paleognathae – a likely consequence of the compromised function of BORIS as a common characteristic across the entire clade. Still, we cannot exclude some functionality in at least some neognathous species. In the human germline and in human cancer cells, 23 alternatively spliced transcripts of BORIS with varying sets of zinc fingers were identified [83]. Therefore, one could expect that mutations leading to a reduced number of zinc fingers, as observed in the Passeriformes, do not necessarily result in a loss of function. However, in species such as *Acanthisitta chloris*, *Amazona aestiva*, or *Apaloderma vittatum*, the presence of premature stop codons or frameshifts in the zinc-finger domain strongly suggests a loss of function. Transcriptome studies could help to decide this question. Searching currently available transcriptomes from six passerine birds and five different tissues each [84], did not reveal evidence that BORIS mRNA is present in these species (results not shown).

The process of gene damage must have started soon after the split of Neognathae and Paleognathae about 100 million years ago and continues to the present. This is supported by our finding that young, species-specific repetitive elements are over-represented within the degrading BORIS loci of neognathous species. Accordingly, we consider the BORIS gene as a gene currently undergoing pseudogenization in Neognathae. Besides mutation and gene duplication, loss of genes through pseudogenization is an important driver of evolution [44, 85–87]. There are numerous examples for lineage-specific events of convergent progressive pseudogenization due to, or followed by, altered selective pressure. While in some cases, the origin for the altered selection is quite obvious [54, 88–91], it is much more obscure or unknown in others [92–94], for instance in BORIS. Taking a closer view on the similarities and differences between the two infra-classes of birds might provide insights.

The taxonomic classification of birds into Neognathae and Paleognathae is based on morphological differences of the Pterygoid-Palatinum Complex (PPC) of the adult skull [97]. In contrast to Paleognathae (∼ 50 known species), Neognathae exhibit a high taxonomic diversity, encompassing approximately 10 000 species, and are characterized by greater cranial kinesis, the movement of skull bones relative to each other, which may have facilitated their extensive diversification as well as their enhanced vocal capabilities [98–100]. In songbirds, it has been shown that the ability of vocal learning is associated with a specific gene expression pattern in the brain [104]. Due to its descent from a transcriptional regulator and the conserved structure of the zinc finger domain, BORIS likely acts as a transcriptional regulator in birds. However, there is currently no indication that BORIS is involved in transcriptional regulation in neuronal tissue.

Beyond differences in cranial morphology, Paleognathae and Neognathae also differ in the morphology of their sex chromosomes. Sex determination in birds relies on ZW chromosomes with heterogamy in females (ZW) and homogamy in males (ZZ). While most Paleognathae have retained ZW homomorphism with extensive and well-conserved pseudoautosomal regions and recombination rates comparable to autosomes, Neognathae display a pronounced ZW heteromorphism with small, gene-poor W chromosomes [101–103]. BORIS is known for its testis-specific expression in mammals [26]. For instance, it has been shown that BORIS regulates other testis-specific genes during spermatogenesis in mice [41, 83, 105]. Therefore, it seems plausible to consider a relationship between the varying characteristics of W chromosomes and the conservation status of BORIS between Paleognathae and Neognathae. However, the origin of BORIS from a duplicate of CTCF does not support a link to sex chromosomes, as both CTCF and BORIS are genuinely autosomal genes.

To further explore a possible involvement of BORIS in the reproductive strategies of birds, we asked whether Neognathae and Paleognathae differ in sperm morphology. We compared the morphological characteristics of sperm in species with and without conserved BORIS. We could show that larger sperm midpiece in neognathous species correlates with altered selective pressure on the BORIS gene compared to Paleognathae and reptiles with a conserved BORIS gene. Genes encoding proteins involved in reproduction are known for their rapid evolution [106–108]. Previous authors suggested post-copulatory sexual selection as the main driver behind this rapid evolution, as reproductive proteins involved in sperm competition tend to evolve specifically fast [109]. This applies particularly to species with promiscuous mating systems [110–112], where sexual selection drives diversity in sperm morphology and function, including variations in the size and structure of the midpiece, which are crucial for motility and successful fertilization [113–117].

Sperm competition and post-copulatory sexual selection are often accompanied by promiscuous mating styles [109–111]. Despite the social monogamy observed during parental care in many neognathous species, genetic analyses of clutches in a variety of bird species have shown that extra-pair paternity is common among Paleognathae and Neognathae [118, 119]. Thus, sperm competition can be assumed to occur throughout the entire avian phylogeny, regardless of whether BORIS is conserved or not, which makes an association between the conservation status of BORIS and sperm competition less likely. Immler et al. [120] could show that the duration of sperm storage in the female reproductive tract also plays a role in the evolution of the sperm. In pheasants, sperm size traits are negatively associated with the duration of sperm storage, indicating that prolonged sperm-female interaction might influence sperm evolution [120].

One notable reproductive difference between Paleognathae and most Neognathae is the presence or absence of external genitalia. While Paleognathae retain external genitalia, most neognathous species have lost this trait [121]. Only three percent of avian species, belonging to two main clades, have retained the ancestral copulatory organ: the Paleognathae and Anseriformes. All other birds have lost the penis-like structure often referred to as intromittend organ (IO) [122–125]. There is a conserved developmental stage of external genital development among all amniotes that suggests a single evolutionary origin of amniote external genitalia [126]. This implies that, in analogy to the presence or absence of the BORIS gene, the presence of an IO is the ancestral state and the loss of an IO is a derived state. The alteration in copulatory anatomy might require different sperm characteristics. Assuming that BORIS exhibits a similar expression pattern and functionality in birds as in mammals, it would be plausible to assume that the reduced selective pressure on BORIS in Neognathae could be functionally related to the altered genital morphology. However, whether there is actually a causal relationship between BORIS degradation, the absence of an IO, and altered sperm morphology remains highly speculative at this point. Clarification can be expected from dedicated expression studies with a focus on reproductive tissues.

## Materials and Methods

### Extraction of genomic DNA

We purchased fresh liver from chicken (*Gallus gallus*) at the local supermarket and cut it into cubes of 2 cm edge length. To avoid contamination with DNA from other poultry, the portions were transferred to 2.8 % sodium hypochlorite solution (NaClO, as in «DanKlorix») for 10 min and washed 3× in PBS (phosphate buffered saline). The pieces were stored in plastic tubes at −20 *^◦^*C. For DNA extraction, the outer layer of the frozen tissue was removed, and samples with a volume of ∼1 mm^3^ were collected from the inside material. After homogenisation in 180 μL lysis buffer, extraction of the genomic DNA was carried out using the Macherey & Nagel NucleoSpin™ Tissue kit. DNA extraction was performed according to the manufacturer’s specifications, with the modification that, after addition of 25 μL Proteinase K, we incubated the sample for 20 min, instead of 1 h to 2 h, at 56 *^◦^*C and 300 rpm. Prior to elution, the silica column was centrifuged at 13 000 *g* for 5 min and dried for another 5 min. The DNA was eluted in 100 μL Tris (10 mm, pH 8.4) and quantified with a NanoDrop™ 2000 spectrophotometer (ThermoFisher Scientific). DNA quality was verified on a 1.0% agarose gel.

### PCR amplification and sequencing

We designed five sets of PCR primers to obtain from chicken genomic DNA five overlapping fragments of a 3.7 kb region targeting the *G. gallus* BORIS locus (see Table 2; Figure 1). To amplify the selected fragments, we used standard PCR conditions: 1. Initial denaturation (94 *^◦^*C, 3 min), 2. Denaturation (94 *^◦^*C, 30 s), 3. Annealing (50 *^◦^*C, 3 s), 4. Elongation (72 *^◦^*C, 3 min), 5. Final extension (72 *^◦^*C, 10 min), with 36 cycles of steps 2–4. The amplified DNA was quantified with a NanoDrop™ 2000 spectrophotometer (ThermoFisher Scientific) and separated on a 1.0 % agarose gel. Amplicons of the expected size were excised from the gel, extracted using the Macherey & Nagel NucleoSpin™ gel and PCR clean-up kit, and sequenced in forward and reverse orientation using the primers displayed in Table 2. Sanger sequencing was performed at Eurofins Genomics, 85560 Ebersberg, Germany. Invdividual sequencing reads were assembled using the phred/phrap/consed package [71, 72]. The resulting contigs were aligned to the *Gallus gallus* reference genome with MUSCLE [73]. Alignments were visualized with SEAVIEW [74].

### Data collection and generation of ORFs

After identifying bird genomic contigs/scaffolds containing BORIS or—if this was unsuccessful—its neighbour genes PCK1 and RBM38 by NCBI BLAST (https://blast.ncbi.nlm.nih.gov/Blast.cgi), using human and bird proteins as queries (BORIS: sp|Q8NI51|CTCFL_HUMAN; PCK1: sp|P05153|PCKGC_CHICK; RBM38: sp|Q5ZJX4|RBM38_CHICK), we downloaded from the NCBI database the respective sequences of 59 bird species under the accessions listed in Supplementary Tables S1 and S2.

The 59 sequences were translated into six open reading frames (ORFs) using the Emboss tool GETORF with parameters «-minsize 90 -find 0» to include all regions between Stop codons of at least 30 AA length [75]. This produced a data set of 5000 to 10 000 ORFs per species, our raw material for the identification and annotation of BORIS fragments and of BORIS neighbor genes in subsequent steps.

### Multiple sequence alignment

Multiple sequence alignments were carried out using the MAFFT v7.304b «einsi» algorithm [76] or MUSCLE [73] with default parameters.

### HMM model and HMM searches

To generate a hidden Markov model (HMM) representative for BORIS from diapsid amniotes (Sauria), to which birds belong, we selected 16 BORIS protein sequences from seven reptilian and nine bird species (Supplementary Table S4). We verified the orthology of these sequences to the protein BORIS in phylogenetic analyses [see 19], generated a multiple sequence alignment from the orthologs using MUSCLE [73], and built a 591 AA BORIS HMM from the alignment using HMMer version 3.1b2 [57]. With the resulting BORIS hidden Markov model, we scanned the ORFs derived from the BORIS syntenic region (spanning genes PCK1 to RBM38; see above) of 59 bird species and annotated ORFs with similarity to BORIS in the corresponding genomic sequence. On the basis of these results, we obtained the status of each individual BORIS zinc finger in each bird species and mapped it onto a consensus bird phylogeny.

### Identification of PCK1 and RBM38 orthologs

Using *Columba livia* PCK1 (tr|A0A2I0M656|A0A2I0M656_COLLI) and RBM38 (tr|A0A2I0M622|A0A2I0M622_COLLI) as queries, we performed BLASTP searches restricted to the taxon «Aves» (Taxonomy ID: 8782) at the NCBI databases (https://blast.ncbi.nlm.nih.gov/Blast.cgi). We downloaded PCK1 blast hits from 76 bird species and RBM38 blast hits from 44 species in fasta format, aligned the corresponding sequences with MUSCLE [73] and visually inspected the alignments using SEAVIEW [74].

### Identification and analysis of repetitive sequences

For comprehensive repeat detection in bird genomes, we combined de novo and library-based identification of repeat elements. For de novo identification, we ran RepeatModeler version 2.0.1 [77] with parameters «-engine rmblast» and «-LTRStruct» on 18 selected bird whole genome sequences (Supplementary Table S3). Typically, several hundred repeat elements per species were detected de novo. We concatenated all repeat elements of the 18 species and eliminated redundancy by clustering with CD-HIT [78] and VSEARCH [79]. Clustering reduced the number of repeat elements from 7372 (overall) to 6666 distinct repeat clusters at 90 % identity threshold. In addition, we identified MITEs (Miniature inverted-repeat transposable elements) in the 18 bird genomes using MITE Tracker [80] with parameter «-mite_max_len 1000». We concatenated the 5988 individual MITEs of 18 species and obtained 4557 MITE clusters with 90 % identity threshold. We then combined the clustered RepeatModeler and MITE Tracker results to construct a comprehensive bird de novo repeat library. We used the combined de novo library and a standard repeat library (Dfam release 3.2; https://dfam.org/home) to mask repeats in the 18 bird genomes in independent RepeatMasker runs. A typical RepeatMasker run contained the following parameters: «-engine rmblast -pa 3 -s -lib $LIB -dir $DATADIR -xsmall -gff -xm -nolow input.fasta». After repeat-masking, we collected for each bird species the repeat content within 100 kb around the BORIS zinc finger region (ZF region start ±50 kb), roughly corresponding to the PCK1 to RBM38 syntenic region. Finally, we analysed in detail repeat contents of the ∼13 kb BORIS consensus locus in selected species using custom scripts and a local JUPYTERLAB instance (python 3.10.9, jupyter server 2.4.0, jupyterlab 3.6.1, pandas 1.5.3, seaborn 0.12.2, and matplotlib 3.7.1).

### Calculation of substitution rates

To investigate the ratio of nonsynonymous vs. synonymous mutations in the BORIS coding region, we first generated a codon-based multiple sequence alignment of BORIS coding sequence from 19 neognathous and 10 paleognathous birds with an intact gene using MACSE (Multiple Alignment of Coding SEquences) release v2.03 [81]. Then we removed poorly aligned positions and divergent regions of the alignments using GBLOCKS Version 0.91b with parameters «-t=c -b2=“(#seq/2)+1+0.5” -b4=2 -b5=a -d=y», with «#seq» specifying the number of sequences in the alignment [82]. Next, we quantified the number of synonymous and nonsynonymous substitutions on each branch using CODEML of the PAML package (parameters: runmode = -2; seqtype = 1; CodonFreq = 2; NSsites = 0) [67]. The tree topology passed to CODEML corresponded to the literature-based consensus phylogeny depicted in Figure 2. Finally, we stored pairwise *dN/dS* ratios from the CODEML output in a table and used a JUPYTER notebook instance (python 3.10.9, jupyter server 2.4.0, jupyterlab 3.6.1, numpy 1.24.2, and matplotlib 3.7.1) for transforming the data into numpy arrays and plotting. The counts of nonsynonymous and synonymous substitutions (*N* × *dN*, *S* × *dS*, respectively) are obtained by multiplying the rates (*dN* and *dS*) with the number of nonsynonymous and synonymous sites (*N* and *S*), as given in the CODEML output table.

### Comparison of sperm morphological data

To compare morphological characteristics of sperm cells between neognathous and paleognathous birds, we extracted measurements on the length of sperm head, mid piece, and flagellum for Aves and Reptilia from the Sperm Tree Database (file «spermtree_01_21_22.xlsx» on https://spermtree.org/database/ [70]). Since the database contains data for only two paleognathous species, we combined Paleognathae (two species) and Reptilia (120 species) into a single dataset and compared it to sperm length measurements from Neognathae (554 species). Using pandas 1.5.3 dataframes, seaborn 0.12.2, and matplotlib 3.7.1 under python 3.10.9, we created boxplots to visualize the data. In addition, we carried out non-parametric Kruskal-Wallis tests to determine if differences in the measurements between Neognathae, Paleognathae, and Reptilia are statistically significant, using the above mentioned python framework and the scipy package (v1.10.1).

## Acknowledgements

This research was funded by the Deutsche Forschungsgemeinschaft (DFG, German Research Foundation)-Projektnummer 268236062-SFB 1211 and supported by the Regional Computing Centre of the University of Cologne (RRZK/ITCC) through access to the HPC system CHEOPS (Cologne High Efficient Operating Platform for Science) on which repeat identification was carried out. We thank Yichen Zheng for insightful comments on an earlier version of this manuscript. Special thanks to the scientific community for distributing genome sequence data to public repositories.

## Dedication

This work is dedicated to the memory of Kamel Jabbari (1966-2020).

