## Supplementary material for "Decay of the CTCF paralog BORIS in neognathous birds": suppl-tables.pdf

### Supplementary tables

**Table S1: List of 59 investigated bird species.** Species information and accession numbers of genomic scaffolds containing the BORIS syntenic region from 59 bird species, part 1. Symbols in column 4 denote if a scaffold includes the syntenic block from genes PCK1 to RBM38 (<>) or is partially complete with only the PCK1 (<) or the RBM38 end (>) present. $: taken from Sackton et al. [127].

| Species | Lineage | Accession | Scaffold |
|---|---|---|---|
| *Acanthisitta chloris* | Neognathae; Passeriformes | NW_019776521.1 | <> |
| *Amazona aestiva* | Neognathae; Psittaciformes | LMAW01000002.1 | <> |
| *Anas platyrhynchos* | Neognathae; Anseriformes | NC_051792.1 | <> |
| *Apaloderma vittatum* | Neognathae; Trogoniformes | NW_009709789.1 | <> |
| *Aptenodytes forsteri* | Neognathae; Sphenisciformes | NW_008794747.1 | <> |
| *Apteryx haastii*[$] | Paleognathae; Apterygiformes | PTFD01000001.1 | <> |
| *Apteryx owenii*[$] | Paleognathae; Apterygiformes | PTFC01000003.1 | <> |
| *Aquila chrysaetos* | Neognathae; Accipitriformes | NW_010972709.1 | <> |
| *Balearica regulorum* | Neognathae; Gruiformes | NW_010747151.1 | <> |
| *Calidris pugnax* | Neognathae; Charadriiformes | NW_015090780.1 | <> |
| *Callipepla squamata* | Neognathae; Galliformes | MCFN01000399.1 | <> |
| *Calypte anna* | Neognathae; Apodiformes | NW_007620763.1 | <> |
| *Caprimulgus carolinensis* | Neognathae; Caprimulgiformes | JMFU01067627.1 | <> |
| *Cariama cristata* | Neognathae; Cariamiformes | NW_009636176.1 | <> |
| *Casuarius casuarius*[$] | Paleognathae; Casuariiformes | PTFA01000086.1 | > |
| *Chaetura pelagica* | Neognathae; Apodiformes | NW_009953486.1 | <> |
| *Colius striatus* | Neognathae; Coliiformes | NW_010701813.1 | <> |
| *Columba livia* | Neognathae; Columbiformes | NW_004973200.1 | <> |
| *Corvus brachyrhynchos* | Neognathae; Passeriformes | NW_008236180.1 | <> |
| *Corvus cornix* | Neognathae; Passeriformes | NW_018113990.1 | <> |
| *Coturnix japonica* | Neognathae; Galliformes | NC_029535.1 | <> |
| *Crypturellus cinnamomeus*[$] | Paleognathae; Tinamiformes | PTEZ01000102.1 | <> |
| *Cuculus canorus* | Neognathae; Cuculiformes | NW_009243347.1 | <> |
| *Cyanistes caeruleus* | Neognathae; Passeriformes | NW_019776521.1 | <> |
| *Dromaius novaehollandiae*[$] | Paleognathae; Casuariiformes | PTEY01000246.1 | < |
| *Egretta garzetta* | Neognathae; Pelecaniformes | NW_009260622.1 | <> |
| *Eudromia elegans*[$] | Paleognathae; Tinamiformes | PTEX01000028.1 | <> |
| *Eurypyga helias* | Neognathae; Gruiformes | JJRO01094590.1 | <> |
| *Falco cherrug* | Neognathae; Falconiformes | NW_004994897.1 | <> |
| *Falco peregrinus* | Neognathae; Falconiformes | NW_004930514.1 | <> |
| *Gallus gallus* | Neognathae; Galliformes | NC_006107.5 | <> |
| *Gavia stellata* | Neognathae; Gaviiformes | NW_009295348.1 | <> |
| *Haliaeetus albicilla* | Neognathae; Accipitriformes | NW_009767762.1 | <> |
| *Haliaeetus leucocephalus* | Neognathae; Accipitriformes | NW_010972709.1 | <> |
| *Lepidothrix coronata* | Neognathae; Passeriformes | NW_016690229.1 | <> |

**Table S2: List of 59 investigated bird species, continued.** Species information and accession numbers of genomic scaffolds containing the BORIS syntenic region from 59 bird species, part 2. Symbols in column 4 denote if a scaffold includes the syntenic block from genes PCK1 to RBM38 (<>) or is partially complete with only the PCK1 (<) or the RBM38 end (>) present. $: taken from Sackton et al. [127].

| **Species** | **Lineage** | **Accession** | **Scaffold** |
|---|---|---|---|
| *Lonchura striata* | Neognathae; Passeriformes | NW_018657563.1 | <> |
| *Melopsittacus undulatus* | Neognathae; Psittaciformes | NC_034427.1 | <> |
| *Merops nubicus* | Neognathae; Coraciiformes | JJRJ01051273.1 | <> |
| *Mesitornis unicolor* | Neognathae; Gruiformes | NW_010159074.1 | <> |
| *Nestor notabilis* | Neognathae; Psittaciformes | NW_009919459.1 | <> |
| *Nipponia nippon* | Neognathae; Pelecaniformes | NW_009000645.1 | <> |
| *Nothoprocta perdicaria*$ | Paleognathae; Tinamiformes | PTEW01000004.1 | > |
| *Numida meleagris* | Neognathae; Galliformes | NC_034427.1 | <> |
| *Opisthocomus hoazin* | Neognathae; Opisthocomiformes | NW_009901057.1 | <> |
| *Parus major* | Neognathae; Passeriformes | NC_031788.1 | <> |
| *Patagioenas fasciata* | Neognathae; Columbiformes | LSYS01003456.1 | <> |
| *Pelecanus crispus* | Neognathae; Pelecaniformes | JJRG01105365.1 | <> |
| *Phaethon lepturus* | Neognathae; Pelecaniformes | NW_010546123.1 | <> |
| *Dryobates pubescens* | Neognathae; Piciformes | NW_009664594.1 | <> |
| *Pseudopodoces humilis* | Neognathae; Passeriformes | NW_005087575.1 | <> |
| *Pterocnemia pennata*$ | Paleognathae; Rheiformes | PTJI01000154.1 | <> |
| *Pygoscelis adeliae* | Neognathae; Sphenisciformes | NW_008825076.1 | <> |
| *Rhea americana*$ | Paleognathae; Rheiformes | PTEV01000005.1 | <> |
| *Serinus canaria* | Neognathae; Passeriformes | NW_007931143.1 | <> |
| *Struthio camelus* | Paleognathae; Struthioniformes | NW_009271896.1 | <> |
| *Sturnus vulgaris* | Neognathae; Passeriformes | NW_014650517.1 | <> |
| *Tauraco erythrolophus* | Neognathae; Musophagiformes | NW_010041494.1 | <> |
| *Tinamus guttatus* | Paleognathae; Tinamiformes | NW_010578138.1 | <> |
| *Tyto alba* | Neognathae; Strigiformes | JJRD01144017.1 | <> |

**Table S3: List of 18 bird species used for repeat analysis.** Species information and NCBI nucleotide accession numbers of 18 genome assemblies used for repeat analysis are shown.

| Species | Lineage | Accession |
|---|---|---|
| *Acanthisitta chloris* | Neognathae; Passeriformes | JAFCHQ000000000 |
| *Amazona aestiva* | Neognathae; Psittaciformes | JAESHV000000000 |
| *Anas platyrhynchos* | Neognathae; Anseriformes | JACEUM000000000 |
| *Balearica regulorum* | Neognathae; Gruiformes | JAAIYC000000000 |
| *Calidris pugnax* | Neognathae; Charadriiformes | LDEH00000000 |
| *Columba livia* | Neognathae; Columbiformes | AKCR00000000 |
| *Cuculus canorus* | Neognathae; Cuculiformes | JAGIYT000000000 |
| *Dryobates pubescens* | Neognathae; Piciformes | JACNMV000000000 |
| *Eudromia elegans* | Paleognathae; Tinamiformes | PTEX00000000 |
| *Gallus gallus* | Neognathae; Galliformes | AADN00000000 |
| *Haliaeetus albicilla* | Neognathae; Accipitriformes | VZSQ00000000 |
| *Melopsittacus undulatus* | Neognathae; Psittaciformes | JAAVWG000000000 |
| *Numida meleagris* | Neognathae; Galliformes | JABXER000000000 |
| *Rhea americana* | Paleognathae; Rheiformes | PTEV00000000 |
| *Streptopelia turtur* | Neognathae; Columbiformes | CABFKC000000000 ? |
| *Struthio camelus* | Paleognathae; Struthioniformes | JJRT00000000 |
| *Theristicus caerulescens* | Neognathae; Pelecaniformes | JAJGSR000000000 |
| *Tyto alba* | Neognathae; Strigiformes | JAEUGV000000000 |

**Table S4: Sequences from diapsid amniotes used to generate the BORIS hidden Markov model.** Origin, NCBI accession number, and length of the nine bird and seven reptilian protein sequences used for HMM generation are shown.

| **Species** | **Lineage** | **Seq-ID** | **[AA]** |
|---|---|---|---|
| | *Birds* | | |
| *Amazona aestiva* | Aves; Psittaciformes | LMAW01000002.1 | 465 |
| *Coturnix japonica* | Aves; Galliformes | XP_015737575.1 | 401 |
| *Empidonax traillii* | Aves; Passeriformes | XP_027737053.1 | 508 |
| *Gavia stellata* | Aves; Gaviiformes | KFV41672.1 | 234 |
| *Lepidothrix coronata* | Aves; Passeriformes | XP_017670709.1 | 451 |
| *Meleagris gallopavo* | Aves; Galliformes | XP_019477849.1 | 612 |
| *Numida meleagris* | Aves; Galliformes | XP_021272595.1 | 545 |
| *Pseudopodoces humilis* | Aves; Passeriformes | XP_014109495.1 | 459 |
| *Serinus canaria* | Aves; Passeriformes | XP_030088730.1 | 446 |
| | *Reptiles* | | |
| *Alligator mississippiensis* | Archelosauria; Crocodylia | KYO42050.1 | 551 |
| *Anolis carolinensis* | Lepidosauria; Squamata | XP_016850798.1 | 581 |
| *Chelonia mydas* | Testudines; Cryptodira | XP_027678081.1 | 665 |
| *Crocodylus porosus* | Archelosauria; Crocodylia | XP_019403720.1 | 510 |
| *Pelodiscus sinensis* | Testudines; Cryptodira | XP_014430204.2 | 664 |
| *Pogona vitticeps* | Lepidosauria; Squamata | XP_020653141.1 | 554 |
| *Python bivittatus* | Lepidosauria; Squamata | XP_025022470.1 | 672 |
